# Widespread adaptive evolution in the photosystems of angiosperms provides new insight into the evolution of photosystem II repair

**DOI:** 10.1101/2024.05.23.595573

**Authors:** Elizabeth HJ Robbins, Steven Kelly

## Abstract

Oxygenic photosynthesis generates the initial energy source which fuels nearly all life on earth. At the heart of the process are the photosystems, pigment binding multi-protein complexes that catalyse the first step of photochemical conversion of light energy into chemical energy. Here, we investigate the molecular evolution at single residue resolution of the plastid-encoded subunits of the photosystems across 773 angiosperm species. We show that despite an extremely high level of conservation, 7% of residues in the photosystems, spanning all photosystem subunits, exhibit hallmarks of adaptive evolution. Through *in silico* modelling of these adaptive substitutions we uncover the impact of these changes on the properties of the photosystems, focussing on their effects on co-factor binding and the formation of inter-subunit interfaces. We further reveal that evolution has repeatedly destabilised the interaction photosystem II and its D1 subunit, thereby reducing the energetic barrier for D1 turn-over and photosystem repair. Together, these results provide new insight into the trajectory of photosystem evolution during the radiation of the angiosperms.

## Introduction

Oxygenic photosynthesis is the process that converts sunlight and carbon dioxide into sugars and oxygen and fuels nearly all life on earth. It is estimated that oxygenic photosynthesis originated ∼3 billion years ago (Planavsky, et al. 2014; Fournier, et al. 2021) and was a landmark event that facilitated the rise of molecular oxygen in the atmosphere and the evolution of aerobic organisms (Eigenbrode and Freeman 2006; Koch and Britton 2008). At the heart of oxygenic photosynthesis are the two photosystem complexes, photosystem I and photosystem II (Blankenship and Hartman 1998). Both photosystems are large multi-subunit protein complexes made up of two parts, the light-harvesting antenna and the reaction centre (Gao, et al. 2018). The function of the antenna complex is to scaffold pigments that absorb and funnel light energy to the reaction centre core (Gao, et al. 2018). The reaction centre uses this light energy for charge separation which ultimately fuels the generation of ATP and NADPH which in turn drive downstream cellular metabolic processes.

The first step of the light-dependent reactions is catalysed by photosystem II, whereby water is oxidised in the thylakoid lumen and the liberated electrons are shuttled through a variety of cofactors to plastoquinone on the stromal side of the thylakoid membrane (Barber 1998). Meanwhile, photosystem I catalyses a later step where reduced plastocyanin from the cytochrome b_6_f complex is oxidised in the thylakoid lumen and the liberated electron is shuttled across the thylakoid membrane to reduce ferredoxin in the stroma (Chitnis 2001). During periods of high light, over-reduction of the internal electron transport chain can result in the formation of reactive oxygen species that damage the photosynthetic apparatus (Hideg, et al. 2002; Asada 2006; Pospíšil 2016). A specific target of this damage is the D1 protein, conferring protective effects on the rest of the photosystem II complex which remains largely undamaged (Mattoo, et al. 1981; Aro, et al. 1993). In particular, the most frequently damaged region of D1 is the ∼40 amino acid long stroma-exposed loop between transmembrane helices D and E, aptly termed the DE-loop (Greenberg, et al. 1987; Aro, et al. 1993; Frankel, et al. 2013). As a result of continuous photodamage, D1 needs to be constantly replaced to maintain photosystem function. Accordingly, D1 has the highest turn-over rate of all photosystem proteins and one of the highest expression levels found in nature (Kettunen, et al. 1997; Li, et al. 2017; Forsythe, et al. 2022).

Eukaryotes gained the capacity for photosynthesis ∼1.5 billion years ago when a single celled eukaryotic organism engulfed a cyanobacterium (Schwartz and Dayhoff 1978; Martin and Kowallik 1999; Hedges, et al. 2004). As the cyanobacterium evolved into the semi-autonomous organelle known as the plastid, its genome underwent a substantial reduction, such that it now harbours <5% of the genes found in its cyanobacterial ancestor (Blanchard and Schmidt 1995; Martin, et al. 2002; Kelly 2021). Despite this large reduction, the gene content and organisation has been highly conserved across the angiosperm lineage (Palmer and Thompson 1982; Jansen, et al. 2007; Zhu, et al. 2016; Robbins and Kelly 2023). Among the ∼80 protein-coding genes that have remained in the plastid genome many encode core photosystem reaction centre proteins. These are single-copy genes, and each has been subject to a low rate of evolution experienced by the plastid genome (Wolfe, et al. 1987; Smith 2015), a direct result of high copy number of the plastid genome and consequently high levels of gene conversion eliminating spontaneous mutations (Birky and Walsh 1992; Khakhlova and Bock 2006). Variation in gene expression, amino acid composition, and genome organisation have further constrained the rate of molecular evolution of genes in the plastid genome (Robbins and Kelly 2023). Together, these factors have resulted in the plastid-encoded photosystem genes being some of the slowest evolving genes found in nature (Cardona, et al. 2019; Oliver, et al. 2021; Oliver, et al. 2023). Given the low rate of molecular evolution it is unknown whether the photosystems have been adaptively evolving during the radiation of the angiosperms, which components of the photosystems have experienced higher or lower rates of adaptive evolution, and how this change has altered the properties of the photosystems themselves.

Here, we present a molecular evolution analysis at the single residue resolution of the plastid-encoded photosystem genes from 773 angiosperm species. We uncover a widespread occurrence of adaptive evolution, namely positive selection and recurrent evolution, in all photosystem genes. Through analysis of the location and type of adaptive changes that occurred, we reveal how these changes have had a significant impact on protein-protein interfaces but not on cofactor binding within the photosystems. Moreover, we reveal a concerted suite of changes that have destabilised photosystem II interactions with the D1 protein thereby reducing the energetic barrier for D1 turn-over and photosystem repair.

## Results

### There is substantial variation in extent of evolution between photosystem subunits

The chloroplast genome in angiosperms contains 20 genes encoding photosystem proteins, 5 from photosystem I (PSI; *psaA*, *psaB*, *psaC*, *psaI* and *psaJ*) and 15 from photosystem II (PSII; *psbA*, *psbB*, *psbC*, *psbD*, *psbE*, *psbF*, *psbH*, *psbI*, *psbJ*, *psbK*, *psbL*, *psbM*, *psbN*, *psbT* and *psbZ*) (Figure 1A). To analyse the molecular evolution of these genes a robust phylogeny was built from a concatenated multiple sequence alignment of all ubiquitously conserved protein-coding sequences from 773 complete angiosperm plastid genomes (see methods). For each photosystem gene, ancestral sequence reconstructions were performed for all 771 internal nodes in the phylogenetic tree, and all non-synonymous substitutions were identified and mapped to branches in the tree.

**Figure 1.**
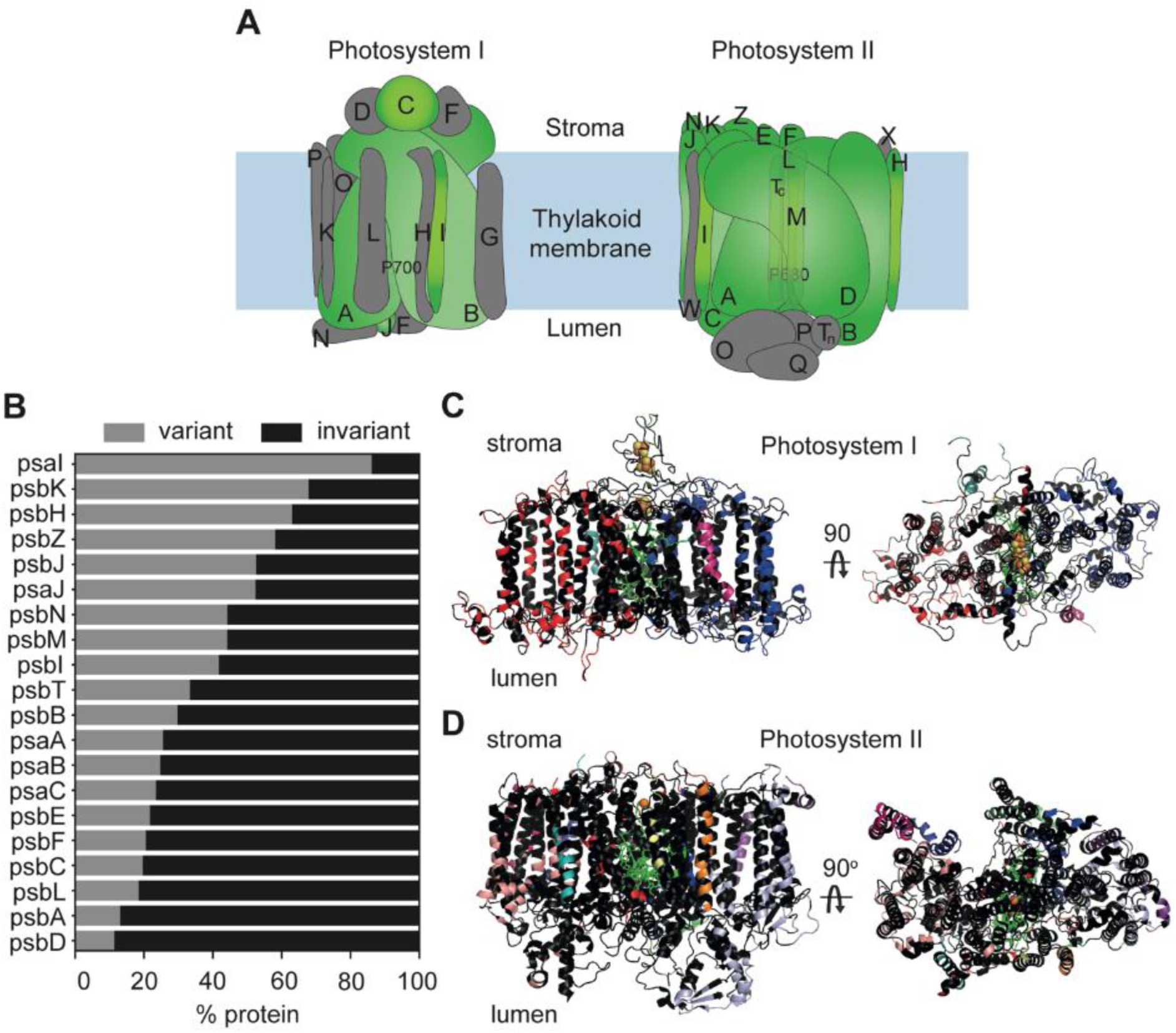
Highly conserved regions of the plastid-encoded photosystem proteins in angiosperms. **A**) Schematics of the photosystem I (left) and photosystem II (right) reaction centres with the proteins encoded by the 20 genes analysed in this study highlighted in green. Proteins in grey were not ubiquitously conserved in the plastid genome among the species analysed. **B**) Bar chart showing the percentage of each protein that was variant (grey) versus invariant (black) in the 773 angiosperm species dataset. Genes are ordered by decreasing percentage of protein that is variable from top to bottom. **C**) Crystal structure of the proteins analysed in photosystem I (PDB: 2O01) with invariant residues highlighted in black. Red, PsaA; blue, PsaB; dark green, PsaC; pink, PsaI and teal, PsaJ. For reference, the electron transfer pathway pigments are shown as green sticks and iron-sulfer centres as orange/yellow spheres. **D**) Crystal structure of the genes analysed in photosystem II (PDB: 7OUI) with invariant residues highlighted in black. Red, PsbA; light blue, PsbB; salmon, PsbC; blue, PsbD; pale green, PsbE; dark green, PsbF; purple, PsbH; teal, psbI; dark blue, PsbK; green, PsbL; orange, PsbM; pale yellow, PsbT and pink, PsbZ. Pigments are shown as green sticks and the Fe^2+^ ion as an orange sphere.

Although 74% (2850/3870) of all residues exhibited no variation (*d*_N_ = 0, Figure 1B - D), 6522amino acid substitutions were identified at the remaining 1020 residues, with substitutions occurring in all 20 photosystem proteins (Figure 2A). Moreover, there was a 17-fold difference between the photosystem protein with the largest and fewest amino acid substitutions per residue across the 773 species phylogeny, corresponding to 11 substitutions per residue for PsaI (encoded by *psaI*) and 0.65 substitutions per residue for D1 (*psbA*), respectively (Figure 2B). Thus, although the plastid-encoded photosystem genes have evolved slowly during angiosperm evolution, evolution has occurred and there is significant variation in evolutionary rate between photosystem components.

**Figure 2.**
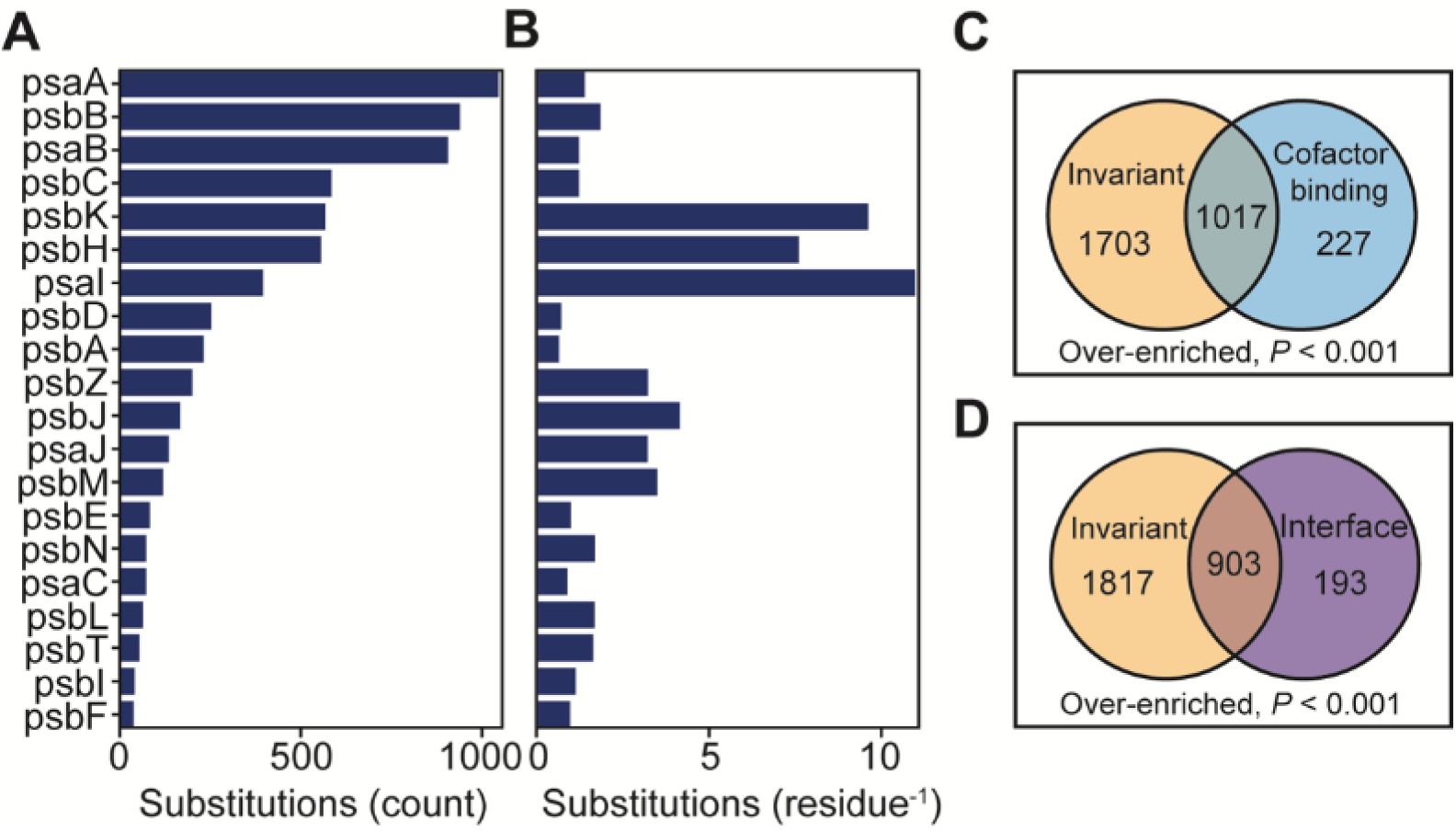
Amino acid substitutions in plastid-encoded photosystem genes during radiation of the angiosperms. **A**) Horizontal bar chart of the raw count of amino acid substitutions inferred to have occurred across the phylogeny for each protein. Proteins are ordered by decreasing number of amino acid substitutions from top to bottom, respectively. **B**) Horizontal bar chart of the substitution count per residue for each protein. Proteins are ordered as in **A**. **C**) Venn diagram showing the overlap between invariant residues (yellow) with cofactor binding residues (blue). **D**) Venn diagram showing the overlap between residues that are invariant (yellow) with those that are at the protein-protein interfaces between subunits (purple). *P*-values shown below Venn diagrams are the results of hypergeometric tests.

### Despite strong functional constraint there is evidence of widespread adaptive evolution in the photosystems of angiosperms

To gain insight into the functional constraints and adaptive evolution of the photosystems in angiosperms we first analysed the sites that exhibited no variation in the 773 species dataset. This revealed that there was an over-enrichment of invariant sites in the cofactor binding residues of the photosystems (hypergeometric test, *P* < 0.001; Figure 2C). Furthermore, there was a significant over-enrichment of invariant sites at subunit interfaces in both photosystems (hypergeometric test, *P* < 0.001; Figure 2D). Thus, the requirement to bind cofactors and form inter-subunit interfaces has resulted in severe constraint on the extent of molecular evolution of the photosystem genes and can explain 58% of all invariant sites in these complexes.

Although sites involved in cofactor binding and subunit interfaces exhibited strong functional constraint, substantial variation is present. This raised the question as to whether this variation exhibits evidence of adaptive evolution, and if so, whether this change has altered the properties of the photosystems. As recurrent evolution and positive selection are widely regarded as signatures of adaptation to common selective constraints (Yang, et al. 2000; Wood, et al. 2005; Kapralov and Filatov 2007; Losos 2011; Stern 2013; Ord and Summers 2015), we sought to determine whether either of these hallmarks could be detected at variable sites in photosystem genes. Given that we had precisely mapped all amino acid substitutions to specific branches in the angiosperm phylogeny, it was possible to directly address the question as to whether any of the amino acid substitutions were recurrent in multiple branches of the phylogeny (a recurrent amino acid substitution is here defined as an amino acid substitution X → Y, where X ≠ Y, that has occurred more than once at a given site in a protein sequence at a frequency greater than expected by chance; see methods). Remarkably, 55% (3620/6522) of all amino acid substitutions were recurrent and unlikely to have evolved by stochastic co-occurrence of random mutations in the absence of selection (*P* < 0.001, Figure 3A). These recurrent substitutions were located at 275 sites across all 20 proteins and comprised 386 unique patterns of amino acid substitution (Figure 3B, Supplementary Table S1). Similarly, there was strong evidence for pervasive positive selection acting at 35 sites in 13 genes (Figure 3C and D, Supplementary Table S2). Moreover, 30 of these 35 positively selected sites were also identified in the set of recurrent substitutions demonstrating a substantial overlap between these two hallmarks of adaptive evolution (hypergeometric test, *P* < 0.001). Thus, there is robust evidence that 280 sites across all 20 plastid encoded photosystem genes have evolved adaptively during the radiation of the angiosperms.

**Figure 3.**
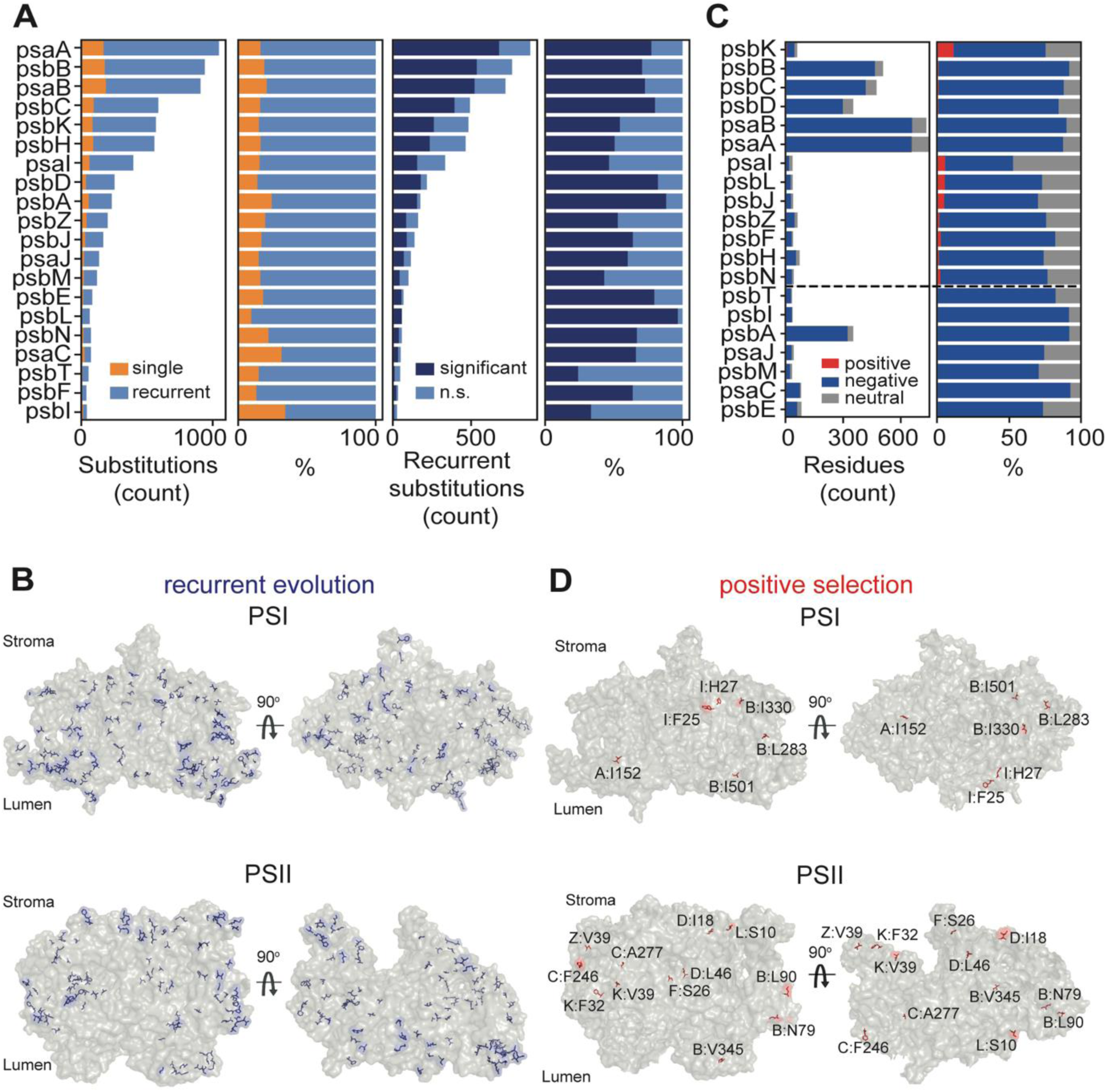
Adaptive evolution in the plastid-encoded photosystem proteins during the radiation of the angiosperms. **A**) Left two bar charts show amino acid substitution counts (left) and percentage of substitutions (right) that are single occurrence (orange) or recurrent (blue), i.e., a specific amino acid replacement that has occurred once or more than once at a given residue, respectively. Proteins are ordered by decreasing number of substitutions inferred to have occurred. Right two bar charts show the proportion of recurrent substitutions that occurred more frequently than expected by chance (*P* < 0.001, dark blue) as compared to those that did not (light blue). Raw counts are shown on the left and percentage of recurrent substitutions shown on the right. **B**) Bar charts showing the number (left) and percentage (right) of residues under positive selection (red), purifying selection (blue) or subject to neither (grey). Proteins are ordered by decreasing number of sites under positive selection from top to bottom. All proteins below the dashed line have no sites under positive selection. **C and D**) Structures of photosystem I (2O01) and photosystem II (7OUI) with modelled sites of recurrent evolution highlighted in blue and sites subject to positive selection shown in red, respectively.

### There is no significant difference in the extent of adaptation between photosystem complexes during angiosperm evolution

We next sought to determine whether adaptive evolution had been more prevalent in either the proteins of photosystem I or photosystem II during the evolution of the angiosperms. Among the 280 adaptively evolving sites, 127 were in photosystem I and 153 were in photosystem II. Given the number of adaptively evolving sites in a protein was strongly correlated to its length (R^2^ = 0.67, *P* < 0.001, Supplementary Figure S1), we compared the number of adaptively evolving sites per residue between subunits of photosystem I versus photosystem II. Although this uncovered a 10-fold difference between the proteins with the highest and lowest levels of adaptive evolution corresponding to 0.250 adaptive sites per residue in PsaI and 0.026 adaptive sites per residue in PsbF, respectively (Figure 4A), there was no significant difference in the level of adaptive evolution between photosystem I and photosystem II subunits (t-test, *P* < 0.05; Figure 4B). Thus, the levels of adaptive evolution experienced by photosystem I and photosystem II has been equivalent during the radiation of the angiosperms.

**Figure 4.**
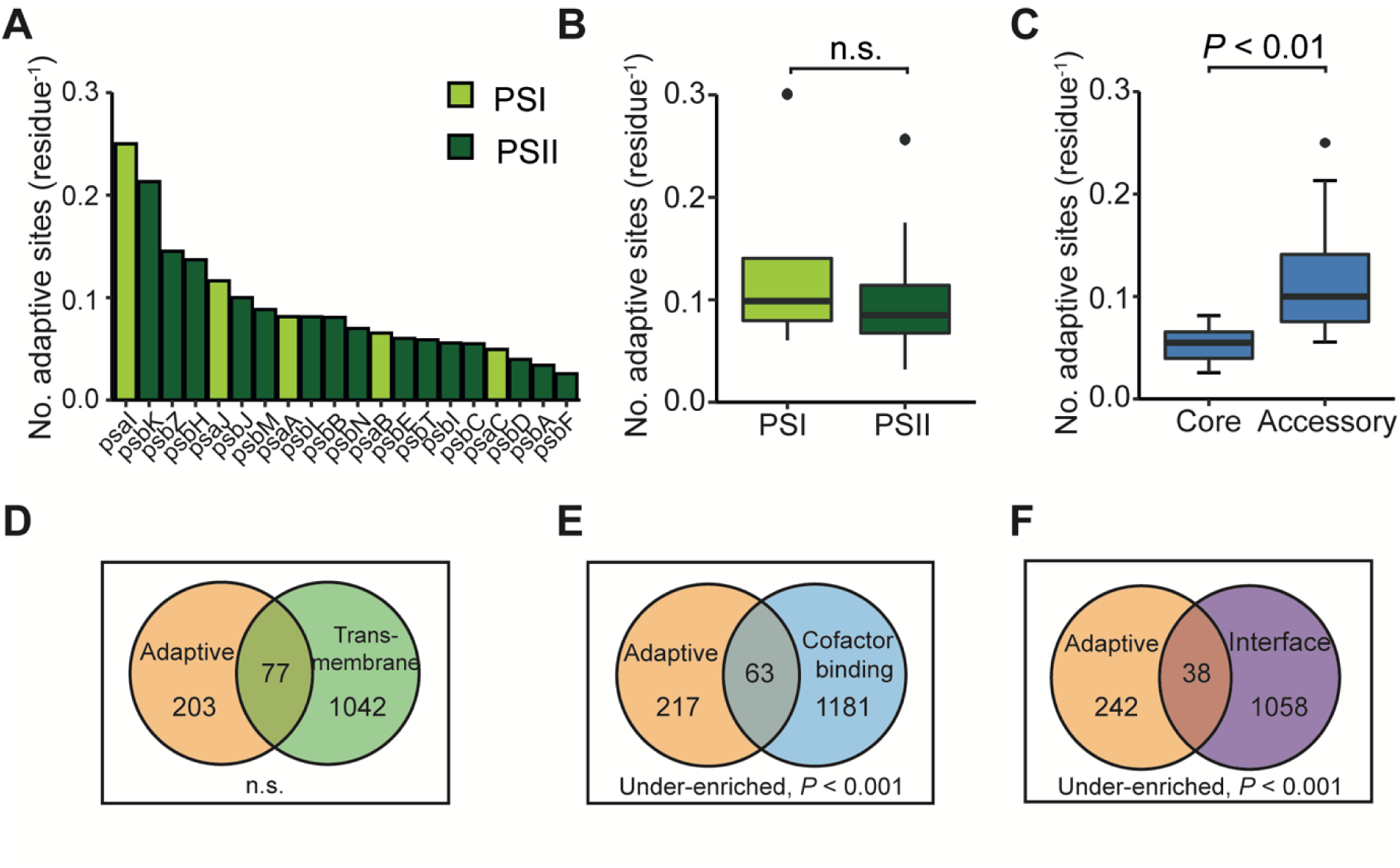
Analysing regional and structural enrichment of adaptive evolution in the photosystems. **A**) Bar chart showing the number of sites under adaptive evolution in the 20 photosystem proteins analysed corrected for protein length in decreasing order from left to right. Proteins in photosystem I (PSI) and photosystem II (PSII) are in light green and dark green, respectively. **B**) Boxplot showing data in (**A**) grouped by photosystem complex. Two-sample t-test, *P* > 0.05. **C**) Boxplot showing data in (**A**) grouped by photosystem region. Core proteins included PsaA, PsaB, PsaC, PsbA, PsbD, PsbC, PsbD, PsbE and PsbF. Peripheral proteins included PsaI, PsaJ, PsbH, PsbI, PsbJ, PsbK, PsbL, PsbM, PsbN, PsbT and PsbZ. *P*-value shown derived from two-sample t-test. **D-F**) Venn diagrams showing the overlap between adaptively evolving sites (orange) and transmembrane residues (green), cofactor binding residues (blue) and inter-subunit residues (purple), respectively. *P*-values shown are the results of hypergeometric tests.

### Adaptative evolution has been limited in regions of the photosystems required for energy and electron transfer

Given that the extent of adaptive evolution was comparable between photosystem complexes, we next sought to investigate whether there was regional enrichment of adaptation within the photosystem structures. Specifically, we wanted to explore whether more adaptation had occurred in the core versus accessory proteins of the photosystem reaction centres, transmembrane versus extramembrane regions or functional regions such as cofactor binding or protein-protein interfaces. To test the former, core proteins were defined as subunits binding the redox cofactors forming the electron transport chain (PsaA, PsaB, PsaC, D1 and D2), internal antenna proteins (PsbB and PsbC) and cytochrome b559 (PsbE and PsbF). Meanwhile, accessory proteins included the remaining 11 low molecular weight (<10 kDa) proteins that are on the periphery of the core reaction centre (PsaI, PsaJ, PsbH, PsbI, PsbJ, PsbK, PsbL, PsbM, PsbN, PsbT and PsbZ). This revealed adaptive sites were significantly more prevalent in the accessory proteins compared to the core proteins of the photosystem reaction centres, with a mean fold difference of 2.2 (t-test, *P* < 0.01; Figure 4C). Meanwhile, there was no significant enrichment of adaptive sites in either the transmembrane or extramembrane regions of the photosystem proteins (Figure 4D). Lastly, adaptive sites were significantly under-enriched in both cofactor binding and inter-subunit interface residues (hypergeometric test, *P* < 0.001; Figure 4E and F), making up 23% and 14% of adaptive sites, respectively. Thus, adaptive evolution has been constrained in regions of high functional constraint including the subunits responsible for energy and electron transfer and residues coordinating cofactors and protein-protein interfaces.

### Adaptive evolution has weakened D1 protein interfaces during angiosperm evolution

Despite being under-enriched at cofactor binding and inter-subunit interface residues, we next sought to assess the impact of the adaptive substitutions that had occurred within these respective regions. We found none of the 63 adaptive substitutions at cofactor binding sites significantly affected ligand binding (Supplementary Table S1). However, of the 38 sites at inter-subunit interfaces 11 substitutions significantly stabilised their respective inter-subunit interface and 12 substitutions significantly destabilised their respective inter-subunit interface (Table 2 and Supplementary Table S1). Of particular note was that all recurrent substitutions at protein-protein interfaces involving the D1 protein (*psbA*) were destabilising (one-sample t-test, *P* < 0.05; Supplementary Figure S2 and Supplementary Table S1), weakening the association of this subunit with the remainder of the photosystem complex. Mapping these sites onto the photosystem II structure revealed that they were primarily located either in, or in contact with, the stromal DE-loop of D1 (*psbA*), a region of known importance for binding and positioning the bicarbonate ion and non-haem iron (Figure 5A). These adaptive changes weakened the interface of the core complex with D1 through a variety of mechanisms. For example, in the DE-loop the recurrent substitution E235A in D1 (*psbA*) results in the loss of polar contacts between E235 in D1 with N264 and W267 in the D2 (*psbD*) protein (Figure 5B). Similarly, although it is not located in the DE-loop region, the L36V substitution in D1 (*psbA*) reduces hydrophobic contacts with F15 and F19 in the transmembrane helix of PsbI (*psbI*) (Figure 5C). Together, this evidence indicates that the photosystems in angiosperms have been adaptively evolving to make it easier to dissociate D1 from the core photosystem complex. This finding is consistent with D1 being the primary recipient of photodamage in photosystem II (Mattoo, et al. 1981; Aro, et al. 1993), having the highest turn-over rate of all subunits in photosystem II (Kettunen, et al. 1997; Li, et al. 2017; Forsythe, et al. 2022), and thus a requirement for continual removal and replacement of D1 to restore photosystem function.

**Figure 5.**
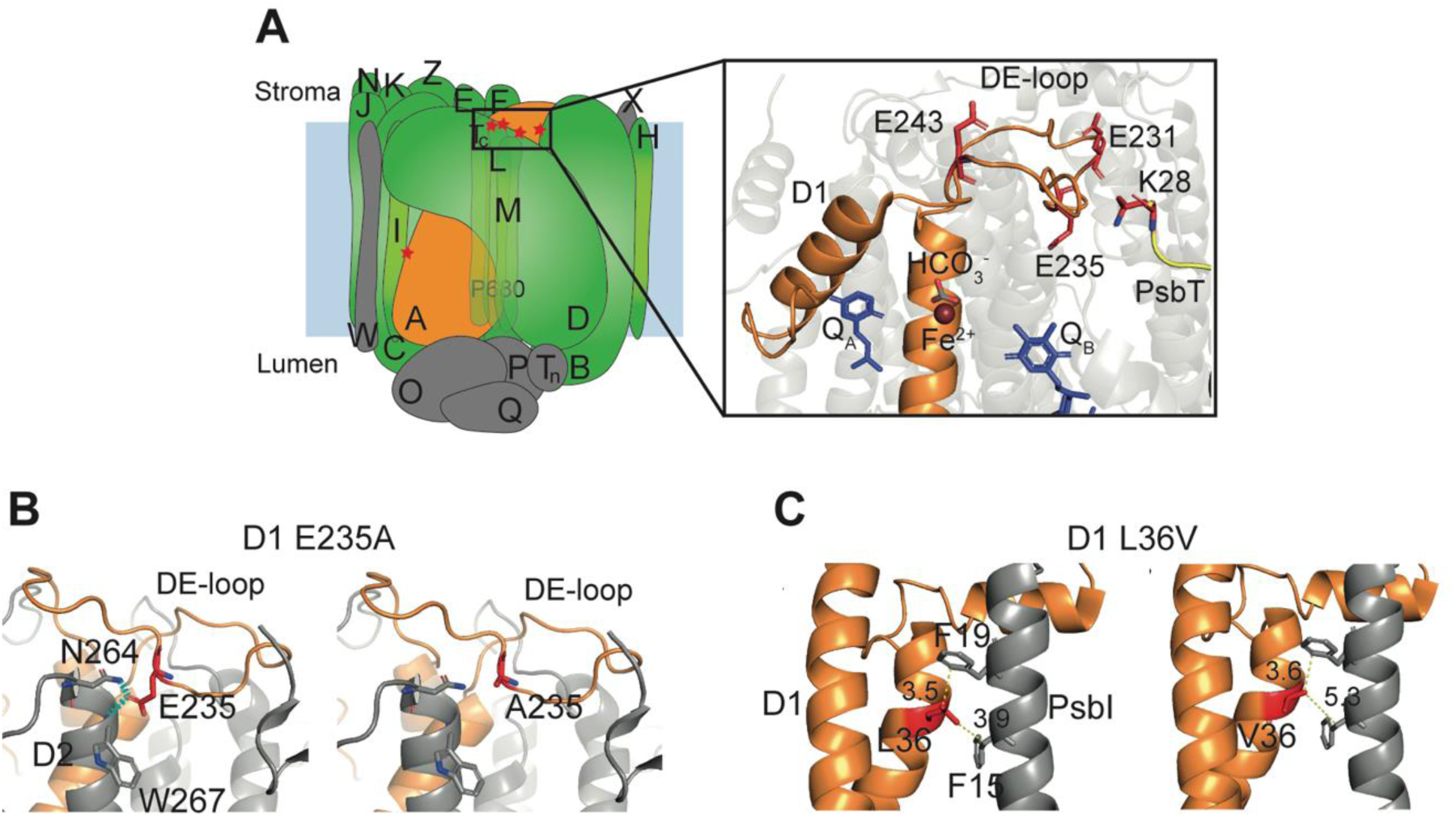
The position and impact of adaptive substitutions on inter-subunit interactions with the D1 (*psbA*) protein. **A**) Schematic mapping the location of substitutions that destabilise inter-subunit interactions with D1. The D1 protein is shown in orange for clarity and sites of substitutions are marked with a red star. A close-up view of the cluster of destabilising substitutions in/around the DE-loop of D1 in the 7OUI photosystem II structure is also provided. Residues at sites of destabilising substitutions are shown as red sticks. For reference, the two plastoquinones (Q_A_ and Q_B_), Fe^2+^ and HCO_3_ ^-^ involved in electron transfer are also shown. **B**) Loss of hydrogen bonds (dashed cyan lines) upon E → A substitution between D1 235 and D2 (*psbD*) N264 and W267. **C**) Reduced hydrophobic interactions upon L → V substitution at 36 in D1 with F15 and F19 in PsbI. Yellow dashed lines show distance between residues given in Å.

## Discussion

Oxygenic photosynthesis underpins nearly all life on earth. At the heart of the process are the photosystems, the molecular machines that facilitate the first step of photochemical conversion of energy from the sun into chemical energy that powers cellular metabolic processes. Since endosymbiosis of the progenitor of the chloroplast in the Archaeplastida lineage (Schwartz and Dayhoff 1978; Martin and Kowallik 1999; Hedges, et al. 2004), the majority of photosystem genes have been retained in the slowly evolving plastid genome resulting in them being some of the slowest evolving genes found in nature (Wolfe, et al. 1987; Smith 2015; Robbins and Kelly 2023). Here, we investigate the molecular evolution of these highly conserved plastid-encoded photosystem genes at single residue resolution over the last 150 million years of angiosperm evolution. We show that hallmarks of adaptive evolution, namely positive selection and recurrent evolution, are widespread in the photosystem subunits of angiosperms. Moreover, we reveal a concerted suite of changes that have destabilised the interaction of photosystem II with its D1 subunit thereby reducing the energetic barrier for D1 turn-over and photosystem repair. Thus, although the photosystems have evolved slowly, there is clear evidence of widespread adaptive evolution during the radiation of the angiosperms.

The analysis presented here focused on angiosperms, and not cyanobacteria or algae, for several reasons. First, the angiosperm clade contains the majority of crop plants, including food plants for direct human consumption, animal feed, biofuels, textiles, building materials and pharmaceuticals (Raskin, et al. 2002; Leff, et al. 2004; Mahapatra, et al. 2021). Consequently, understanding the evolution of their photosynthetic apparatus, i.e., what evolution has tried and tested, holds most potential to inform future crop improvement strategies. Second, a large number of angiosperm plastid genomes have been sequenced allowing for powerful phylogenetic analyses (Christenhusz and Byng 2016). Third, the high degree of gene conservation in the plastid genome of angiosperms enabled a robust phylogenetic tree to be constructed from which adaptive evolution was inferred (Robbins and Kelly 2023). The analysis presented here will provide a foundation and point of comparison for investigation of adaptive evolution in photosystems of other lineages, such as algae or cyanobacteria, in future studies.

The finding that all adaptive substitutions at the inter-subunit interfaces with D1 (*psbA*) destabilise the protein-protein interactions is consistent with extremely high turnover of the subunit (Kettunen, et al. 1997; Li, et al. 2017; Forsythe, et al. 2022), and the concomitant requirement to constantly remove and replace damaged D1 protein. Four of the five of these substitutions were found to be clustered in or around the stroma-exposed DE-loop in D1 (D1: E231Q, E235A, E243G and PsbT: K28T), which has been attributed two main functions. The first is that it provides two ligands (E244 and Y246) that indirectly coordinate the non-haem iron positioned between Q_A_ and Q_B_ *via* positioning a bicarbonate ion (Gao, et al. 2018). Secondly, the region between residues 238-248 in the DE-loop is a site of cleavage during D1 degradation, yielding the 23 kDa fragment (Greenberg, et al. 1987). Moreover, it has been postulated that a damage induced increase in mobility of the DE-loop may signal photosystem II repair (Forsman 2021). In this context it interesting to note that three of the adaptive substitutions found to destabilise a D1 inter-subunit interface were located in the DE-loop region of D1, E231Q, E235A and E243G. Interestingly, it was shown that mutagenesis at two of these sites (E231D and E243Q) did not affect electron transfer but did substantially reduce the half-life of D1 (Mulo, et al. 1997). Given that electron transfer was not affected, the bicarbonate ion and non-haem iron must have still been successfully bound to the photosystem, suggesting the DE-loop has maintained a mostly correct conformation. If this is the case, the reduced half-life cannot be the result of an aberrant DE-loop conformation signalling immediate photosystem II degradation. An alternative hypothesis is that these substitutions result in less photodamage being required to initiate removal and replacement of D1, due to a reduction in the energy required to remove D1 from photosystem II. It is most parsimonious to assume that the adaptive substitutions we uncovered here act in a similar manner.

We also identified a further adaptive substitution, PsbT K28T, which is located adjacent to the DE-loop of D1 and predicted in this analysis to weaken the D1:PsbT protein-protein interface. Importantly, this adaptive substitution should not disrupt important hydrophobic interactions that occur between the neighbouring residues in PsbT (P27 and I29) with F239 in the DE-loop of D1. Without these hydrophobic interactions, the DE-loop has increased flexibility that leads to bicarbonate dissociation and slowed electron transfer between Q_A_ ^-^ and Q_B_ (Forsman and Eaton-Rye 2021). This increased flexibility and resultant aberrant conformation of the DE-loop is thought to be the cause of an increase in turnover of D1 (Forsman and Eaton-Rye 2021). Thus, taken together the findings presented here indicate that the K28T substitution in PsbT is adaptive as it weakens the connection between PsbT and D1 without disrupting the DE-loop conformation, thereby enhancing D1 repair signalling in response to photodamage.

In this analysis we identified 7% of sites in the plastid-encoded photosystem proteins of angiosperms show hallmarks of adaptive evolution. Furthermore, we showed that many of these sites are located at regions of known functionality, with 23% of adaptively evolving sites involved in cofactor binding and 14% of sites at inter-subunit interfaces. However, while we provide biological explanations for substitutions at several adaptively evolving sites, the majority remain unexplained. For some of these unexplained sites, *in silico* prediction of the effect of substitution is impossible due to their location in regions that are not modelled in the photosystem structures. Meanwhile, for the currently unexplained sites that are resolved in the photosystem structures their beneficial trait is not obvious from analysing the change to protein structure. Nonetheless, this study identifies several regions of the photosystems with robust evidence of adaptive evolution and further experimental investigation is required to elucidate the biological importance of these changes.

Our findings show that despite evolving very slowly, that all plastid-encoded photosystem proteins have experienced adaptive evolution during the radiation of the angiosperms. By uncovering these past adaptive trajectories, and identifying key sites that have been altered by natural selection, these findings provide substantial new insight into the present and future evolution of the photosystems in angiosperms.

## Methods

### Phylogenetic tree inference

The plastid genome dataset and phylogenetic tree used in this study was the same as in (Robbins and Kelly 2023). In brief, the full set of sequenced plastid genomes was downloaded from the National Center of Biotechnology and Information (NCBI) in July 2021. Genomes were filtered so that those that did not contain the full complement of canonical plastid genes were removed, resulting in a dataset containing nucleotide, and corresponding protein sequences, for 69 plastid genes for 773 angiosperm species. For each of the 69 genes, a protein multiple sequence alignment was generated using MAFFT L-INS-I (Katoh and Standley 2013) and was used to guide an accurate nucleotide alignment using Pal2Nal (Suyama, et al. 2006). The nucleotide alignments were then trimmed to remove columns containing >90% gaps (Supplementary File S1) and concatenated together and used to identify the best-fit model of nucleotide evolution using IQ-TREE’s inbuilt ModelFinder (Kalyaanamoorthy, et al. 2017). The best-fitting model of nucleotide evolution identified was GTR+F+R7. IQ-TREEs ultrafast bootstrapping method with 1000 replicates was then used to infer a maximum likelihood phylogenetic tree using the concatenated nucleotide sequence and GTR+F+R7 model of evolution (Hoang, et al. 2018). The tree was than manually rooted on the Nymphaeles clade, as they are the most basal angiosperm lineage present in the dataset.

### Inferring non-synonymous substitutions and calculating the rate of non-synonymous substitution

Among the 69 ubiquitously conserved genes above were 20 genes that encode proteins that participate in the photosystem reaction centres, 5 for photosystem I (*psaA*, *psaB*, *psaC*, *psaI* and *psaJ*) and 15 for photosystem II (*psbA*, *psbB*, *psbC*, *psbD*, *psbE*, *psbF*, *psbH*, *psbI*, *psbJ*, *psbK*, *psbL*, *psbM*, *psbN*, *psbT*, *psbZ*). Ancestral state reconstructions were generated for each of the 20 photosystem genes for every node in the maximum likelihood phylogenetic tree using the GTR+F+R7 model of evolution in IQ-TREE (Nguyen, et al. 2014). Non-synonymous substitutions at every internal branch in the tree were inferred by assessing all child-parent relationships in the phylogeny, excluding the Nymphaeles outgroup. These were translated into amino acid replacements at every residue in the protein alignment. Substitutions were tabulated in mutation matrices where columns represent the parent residue identity and rows represent the child residue. Site-wise rates of non-synonymous mutation (*d*_N_) were determined using the FEL method from HYPHY’s software suite (Kosakovsky Pond and Frost 2005).

### Identifying cofactor binding and protein-protein interface residues

Residues involved in cofactor binding and protein-protein interactions were identified using the 2O01 photosystem I (Amunts, et al. 2007) and 7OUI photosystem II (Graça, et al. 2021) structures. To identify cofactor binding residues, all pigments and cofactors associated with the 20 subunits analysed were identified in the photosystem structures. For photosystem I this included 93 chlorophyll a molecules, 2 plastoquinones, 3 beta-carotenes and 3 iron-sulfer centres. Meanwhile, for photosystem II this included 35 chlorophyll a molecules, 4 chlorophyll b molecules, 2 Ca^2+^, 1 Cl^-^, 11 beta-carotene, 1 HCO_3_ ^-^, 4 digalactosyl diacyl glycerol, 6 1,2-distearoyl-monogalactosyl-diglycerise, 2 pheophytin, 2 plastoquinone and 4 sulfaquinovosyldiacylglycerol molecules. Residues coordinating these pigments and cofactors were identified by conducting a search for residues with at least one heavy atom <4Å from an atom in a pigment or cofactor using the NeighbourSearch function in the Bio.PDB package (Hamelryck and Manderick 2003) (Supplementary Figure S3 and Supplementary Table S3). Protein-protein interface residues were identified using the EMPL-EBI PDBsum database (Supplementary Figure S4) (Laskowski, et al. 2018) (Supplementary Table S4).

### Recurrent evolution and positive selection

Recurrent substitutions were defined as amino acid substitutions, i.e., X -> Y, where X ≠ Y, that were inferred to have occurred more than once at a specific residue in the phylogeny. Given that recurrent substitutions could have arisen from chance a Monte Carlo approach was developed to assess whether they were likely to occur by chance given the inferred model of sequence evolution and the phylogenetic relationships between species encapsulated by the tree. To do this, the best-fitting model of protein sequence evolution was identified from the concatenated alignment of all 69 conserved plastid protein sequences and the species tree using IQ-TREE’s inbuilt ModelFinder (Kalyaanamoorthy, et al. 2017). The best-fitting model identified was JTT+F+R7. Branch lengths for protein trees that were constrained to the topology of the species tree were then inferred for each of the 20 photosystem protein alignments using the JTT+F+R7 model of protein evolution. Protein sequences for extant species were then simulated for each of the 20 photosystem genes from the protein tree and ancestral node sequence using IQ-TREE’s alignment simulator, with 1000 replicates. For each of the simulated alignments, ancestral sequences were inferred, and amino acid substitutions counted as described above. Only the recurrent substitutions that had not occurred at the same or higher frequency in any of the 1000 simulations of protein evolution were deemed significant and a hallmark of adaptive evolution.

Sites under positive selection were identified using the FUBAR method from the HYPHY software package (Murrell, et al. 2013). Sites with a posterior probability that the rate of non-synonymous substitution is greater than the rate of synonymous substitution (β > α) of greater than 0.9 were defined as under positive selection.

### Modelling the effect of amino acid substitutions on cofactor binding and protein-protein interfaces

Single amino acid substitutions were introduced into the photosystem I (2O01) and photosystem II (7OUI) structures using PyMol. To estimate the effect of a substitution on cofactor binding, the change in Gibb’s free energy of binding (ΔG_binding_) between protein and ligand was calculated for the wild-type and mutated structures using the PRODIGY-LIG webserver (Vangone, et al. 2018). From these, the change in ΔG_binding_ between the mutated and wild-type structure (ΔΔG_binding_) can be calculated, whereby a negative value indicates an increased binding affinity, and *vice versa*. Similarly, to estimate the effect of substitution at a protein-protein interface, the change in Gibb’s free energy of interaction (ΔG_PPI_) was calculated using BAlaS for both the wild-type and mutated protein structures (Wood, et al. 2020). As before, the change in ΔG_PPI_ upon mutation (ΔΔG_PPI_) was then calculated. For both ΔΔG_binding_ and ΔΔG_PPI_ only values <-0.5kcal.mol-1 or >0.5kcal.mol-1 were considered to substantially alter cofactor binding or protein-protein interactions, respectively (Benevenuta, et al. 2022).

## Data Availability

All data is provided in the supplemental material.

## Author Contributions

ER and SK conceived the study. ER conducted the analysis. ER and SK wrote the manuscript.

---

**Table 1.** Adaptive substitutions that substantially affect protein-protein interface stability (|ΔΔG_PPI_| ≥ 0.5 kcal.mol^-1^).

| Protein | Mutation | Protein-protein interface | $\Delta\Delta G_{\text{PPI}}$ (kcal.mol<sup>-1</sup>) |
| --- | --- | --- | --- |
| PsaB | M640T | B:A | -6.15 |
| PsbF | S26F | F:E | -2.7 |
| PsaI | S25F | I:L | -1.67 |
| PsbE | D68N | D:E:U | -1.53 |
| PsaC | H71P | C:B | -1 |
| PsbE | S72P | E:D | -0.66 |
| PsaI | H27Y | I:B | -0.6 |
| PsaA | D49E | A:E | -0.58 |
| PsbL | N10S | B:L:M | -0.57 |
| PsaJ | I4L | J:F | -0.54 |
| PsbE | C124T | E:H | -0.53 |
| PsbL | S10N | B:L:M | 0.57 |
| PsaI | N3S | I:H | 0.63 |
| PsbD | I18T | D:X | 0.66 |
| PsbE | P72S | E:D | 0.66 |
| PsbA | L36V | A:I | 0.95 |
| PsbT | K28T | A:T:D | 1.36 |
| PsbA | E235A | A:D | 1.43 |
| PsaC | W70S | C:B | 2.37 |
| PsbA | E231Q | A:B | 2.43 |
| PsaC | W70G | C:B | 2.61 |
| PsbB | A471S | B:D | 4 |
| PsaC | H71N | C:B | 4.23 |

