## Supplemental Figures for "Widespread adaptive evolution in the photosystems of angiosperms provides new insight into the evolution of photosystem II repair"

### ***Supplementary Figure S1***

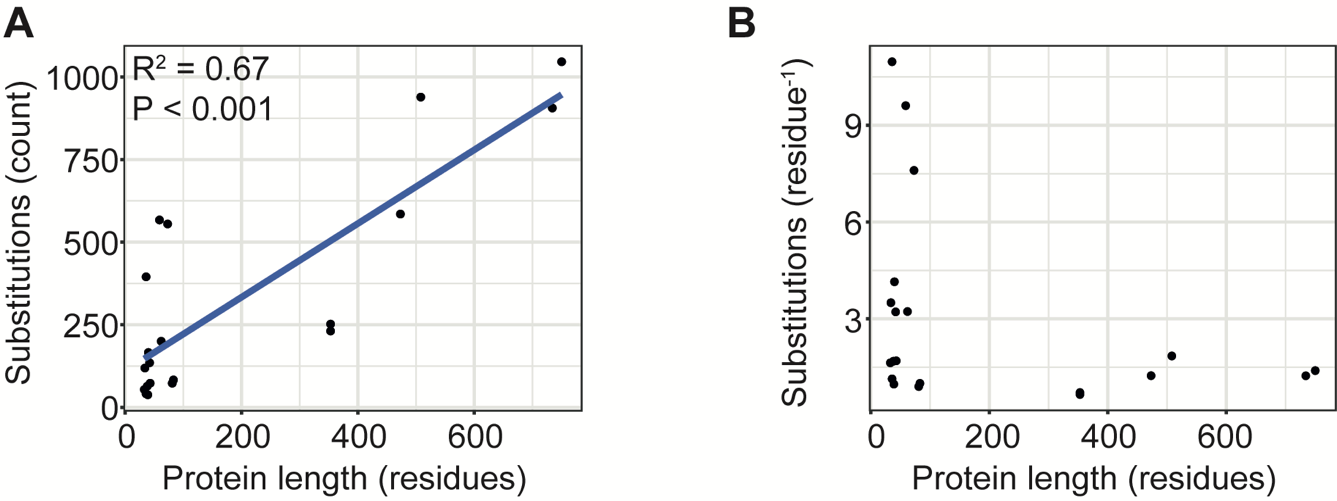

**Supplementary Figure S1**. Relationships between the number of inferred non-synonymous substitutions with protein length during the evolution of our angiosperm phylogeny. **A**) Scatter plot of the non-synonymous substitution count per gene with the length of the protein given in residues. A linear regression line is shown in blue with the associated R^2^ and *P* values given. **B**) Scatter plot of the non-synonymous substitutions per residue versus protein length.

***Supplementary Figure S2***

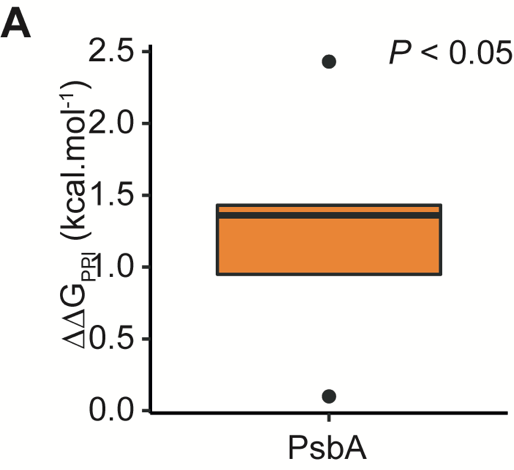

**Supplementary Figure S2**. Boxplot showing the change in inter-subunit interaction energy upon adaptive substitutions at a PsbA (D1) interface. *P*-value shows the result of a one-sample t-test, n=5 (PsbA L36V, E231Q, E235A, E243G; PsbT K28T).

***Supplementary Figure S3***

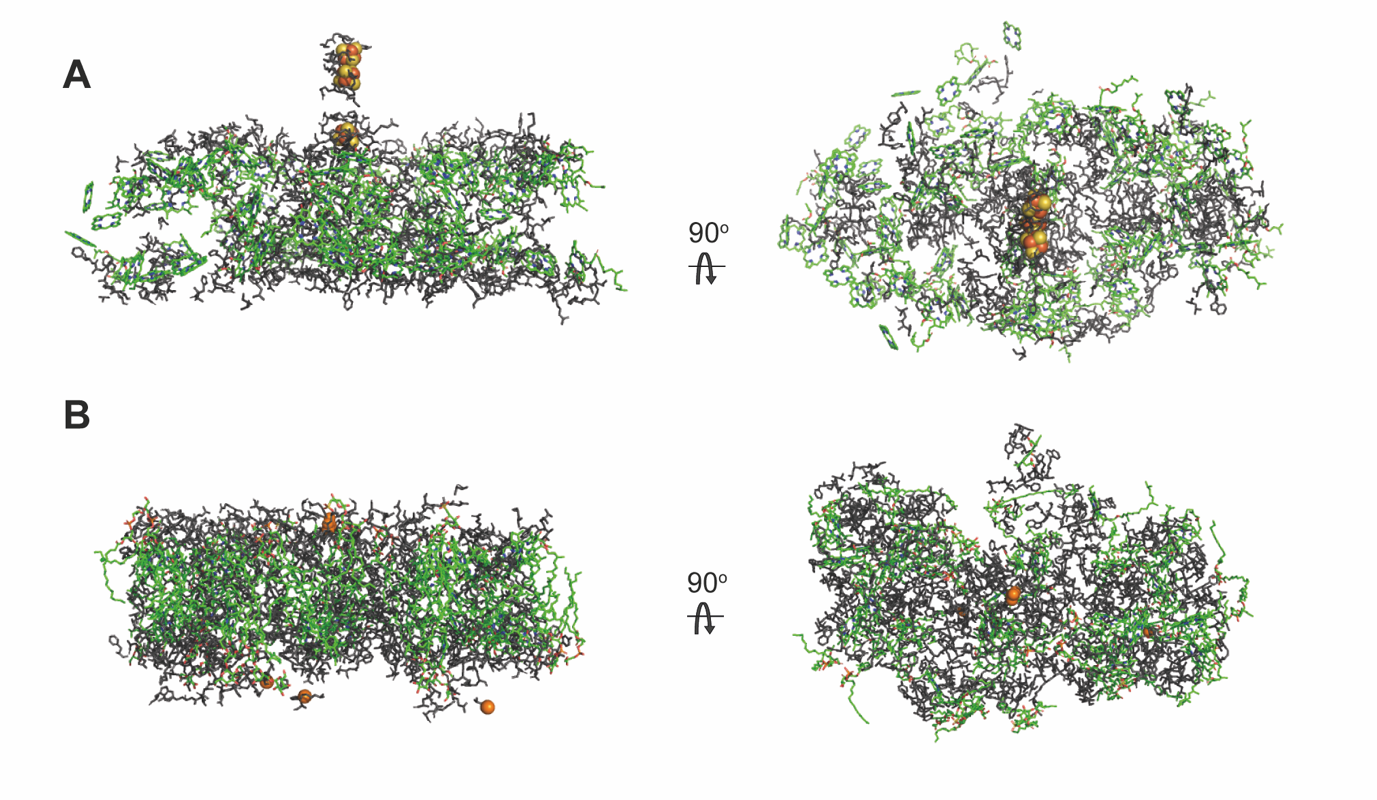
**Supplementary Figure S3.** Cofactor interacting residues in the conserved plastid-encoded photosystem genes. **A**) Analysis of the 2O01 structure of photosystem I. Cofactors associated with psaA, psaB, psaC, psaI or psaJ are shown in green. The 525 residues in these proteins that have an atom within 4Å of a cofactor atom are shown in grey. **B**) Analysis of the 7OUI structure of photosystem II. Cofactors associated with psbA, psbB, psbC, psbD, psbE, psbF, psbH, psbI, psbK, psbL, psbM, psbT or psbZ are shown in green or as orange spheres for small molecules and atoms. The 719 residues in these proteins that have an atom within 4Å of a cofactor atom are shown in grey.

***Supplementary Figure S4***

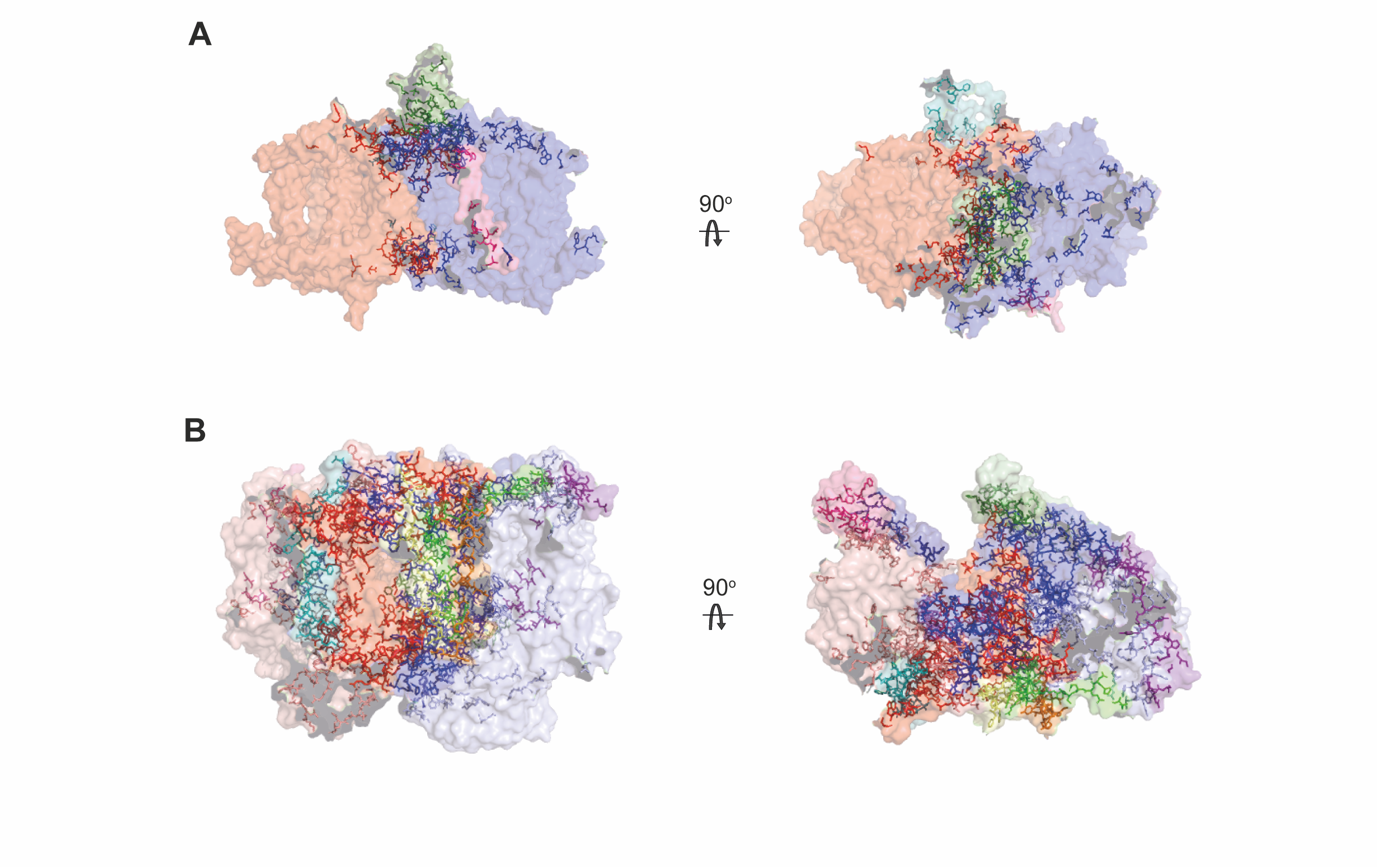

**Supplementary Figure S4**. Inter-subunit interface residues in the conserved plastid-encoded photosystem genes. Only proteins analysed in this study are shown. Residues identified at inter-subunit interfaces shown as sticks. **A**) Analysis of the 2O01 structure of photosystem I. Red, psaA; blue, psaB; green, psaC; pink, psaI and teal, psaJ. **B**) Analysis of the 7OUI structure of photosystem II. Red, psbA; light blue, psbB; salmon, psbC; blue, psbD; pale green, psbE; dark green, psbF; purple, psbH; teal, psbI; dark blue, psbK; green, psbL; orange, psbM; pale yellow, psbT and pink, psbZ.
